# Optical Windows for Transcranial Brain Imaging in Living Mice: Skull Thinning, Clearing, and Beyond

**DOI:** 10.64898/2026.03.08.710408

**Authors:** Yiming Fu, Gewei Yan, Zhentao She, Yingzhu He, Kangqi Liu, Jianan Qu

## Abstract

Longitudinal, noninvasive *in vivo* imaging is essential for studying brain physiology and pathological mechanisms. Advances in transcranial optical windows, including thinned-skull and optical clearing techniques, have markedly improved imaging depth and resolution when combined with multiphoton microscopy. However, their optical performance often deteriorates rapidly, and quantitative studies on long-term stability and the causes of image quality loss remain limited. In this work, we systematically investigated current transcranial window approaches using multiphoton excited fluorescence microscopy (MPEFM) and adaptive optics, examining longevity, optical aberrations, and imaging resolution. Our results reveal that progressive skull regrowth is a fundamental limitation across all window types, leading to substantial declines in signal quality and resolution for conventional MPEFM thereby reducing achievable high-resolution imaging depth. To address these challenges, we developed a localized glucocorticoid (GC) delivery strategy that significantly extends window performance for up to one month. Furthermore, we demonstrated that a GC-loaded hydrogel sealing method effectively suppressed skull regrowth while preserving optimal optical properties, offering a potential and practical route to chronic, high-fidelity transcranial imaging. These findings provide mechanistic insight into window degradation and establish a framework for sustained, long-term *in vivo* brain imaging.

## Introduction

Multiphoton microscopy (MPM) has become a central tool in neuroscience for visualizing fine cellular structures and dynamic cell-cell interactions deep within the living brain^1–5^. Its ability to perform repeated, noninvasive *in vivo* imaging enables detailed investigations of neural function, disease progression, and treatment effects over time^6–11^. However, image quality is often compromised by optical scattering and aberrations introduced by the skull’s opaque and turbid structure, limiting both resolution and penetration depth. The open-skull cranial window is the most used method, allowing for longitudinal high-quality imaging^12,13^. However, it is invasive and can lead to complications such as inflammatory responses, glial activation, permanent alterations in neural connectivity and disruption of cerebrospinal fluid dynamics^12–14^.

Several optical windows have been proposed and developed to address the issues. The thinned skull window minimizes disruption to the brain by removing only the outer surface layer of the skull while preserving a thin underlying layer^15^. To extend imaging duration and field of view (FOV), the polished and reinforced thinned skull (PoRTS) window has been introduced ^16^.This method seals the thinned bone with transparent cyanoacrylate cement, enabling imaging for up to three months with an extended FOV of 3×3 mm. The optical clearing (OC) window achieves optical transparency without thinning the skull by chemically removing lipids, collagens and match the skull refractive index (RI), thereby reducing scattering and enhancing transparency^17,18^. The OC window allows for even larger imaging areas while maintaining mechanical stability, which is essential for experiments that require preserving the skull’s integrity as a reservoir for myeloid cells in the meninges^19,20^. While these non-invasive windows represent significant advancements in brain imaging, few studies have quantitatively evaluated their performance in terms of penetration depth, optical aberrations, and imaging resolution. Moreover, there is limited research on window longevity, which is essential for long-term *in vivo* experiments requiring longitudinal observations over weeks to months. This lack of detailed information hinders researchers from selecting the most appropriate optical window for *in vivo* brain imaging.

In this study, we comprehensively evaluated all four transcranial windows: thinned skull^15^, PoRTS^16^, OC window^17^, and a newly developed thinned-clearing (TC) window in terms of imaging duration, optical aberration, and their capacity for high-resolution imaging using both conventional two-photon excited fluorescence microscopy (TPEFM) and advanced adaptive optics (AO)-TPEFM^4,21,22^. We demonstrated that the TC window is an optimized approach that combines skull thinning with optical clearing, significantly shortening the clearing procedure, particularly in aged mice with vessels embedded in spongy bone, and improving brain fluorescence signal intensity compared with either thinned-skull or optical-clearing windows alone. Through longitudinal imaging, we found that skull regrowth exists in all four types of windows and results in the quick degradation of imaging quality after one week. Using the AO-TPEFM technology^4^, we analyzed optical aberrations across window types and precisely characterized how aberration affects imaging resolution and depth. Recognizing skull regrowth as the fundamental limitation to sustained long-term window performance, we devised methods to suppress regrowth and extend window longevity. Glucocorticoids (GCs), known to impair bone regrowth by decreasing osteoblast numbers and inhibiting bone matrix synthesis while increasing osteoclast numbers and promoting bone resorption^23–25^, were locally applied to the thinned skull, successfully inhibiting regrowth for up to one month. We carefully controlled GC concentration to minimize brain immune responses and found that fluorescence signal remained stable during skull regrowth suppression, highlighting the importance of this intervention for high-quality long-term transcranial imaging. Finally, we developed a transcranial optical window sealed with a GC-loaded transparent hydrogel, enabling repeatable long-term imaging while maintaining high imaging quality over several weeks.

## Results

### Transcranial windows for multiphoton microscopy of the living mouse brain

While various transcranial windows were reported over a decade ago, there has been limited quantitative evaluation of imaging quality and stability over time across different window types. In this work, we first investigated the optical properties of these transcranial windows using conventional multiphoton fluorescence microscopy. We reproduced the thinned skull window, PoRTS window, and OC window following previously reported preparation protocols^15–17^. *Cx3cr1^GFP/+^* transgenic mice were used because their stable GFP expression in microglia under physiological conditions allows both monitoring of potential inflammation caused by window preparation and providing a consistent fluorescence source for evaluating imaging depth and resolution^26,27^.

To quantify and compare the imaging quality over time, we first proposed a uniform procedure for data collection and image formation. Specifically, for each mouse, we acquired fluorescence images of microglial GFP up to 350 µm below the pia using a standard TPEFM with an excitation wavelength of 920 nm (See Methods). To compensate for the attenuation of fluorescence signal at depths, we empirically selected four excitation power levels for different brain depths (Fig. S1a): 10 mW (0–50 µm), 20 mW (50–150 µm), 40 mW (150–250 µm), and 80 mW (250–350 µm). Then the fluorescence intensity was first normalized to the excitation power at each depth range to account for sudden variations in GFP intensity introduced by power adjustments (Fig. S1b). We then plotted the logarithm of GFP intensity versus depth and fitted the data with a linear function *I*=*α* · *d* + *I*_0_ (Fig. 1a, Fig. S1d)^28^. The intercept of the fitted linear function, *I*_0_, was used to quantitatively represent the global fluorescence signal intensity from the entire imaging volume. To improve visualization of deep-brain signals, we further normalized for the exponential decay of signal with depth for display (Fig. S1c). Using this uniform image formation approach, the performance of different windows can be quantitatively compared based on their global signal intensity *I*_0_. Representative results of thinned skull, PoRTS, OC and TC windows are shown in the Supplementary Information (Fig. S1a-d). However, it must be emphasized that the performance of individual windows varies due to many factors, such as surgical preparation, thickness of skull, GFP expression level and mouse age.

**Figure 1:**
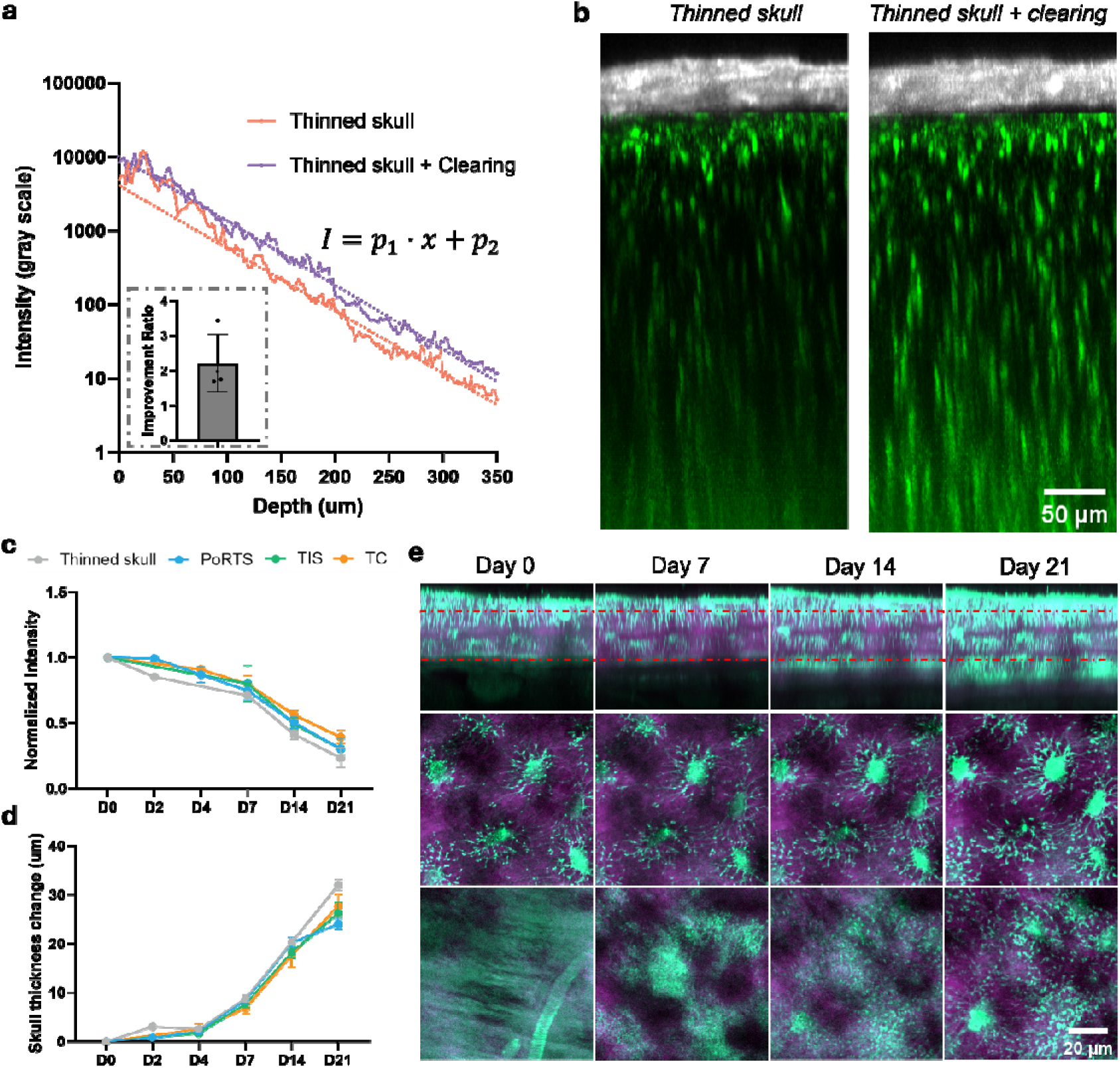
Quantitative characterization of various optical transcranial windows. **a.** Plot of GFP fluorescence intensity versus imaging depth of thinned skull window TC window of the same mouse. Sub-figure shows the mean fluorescence intensity ratio between the TC window and the thinned-skull window. **b.** Maximum projection image of thinned skull window (left) and TC window (right) along y-axis on a *Cx3cr1^GFP/+^* mouse. SHG signal (gray): skull; GFP (green): microglia. **c-d.** Statistics of normalized GFP intensity changes (c) and skull thickness changes (d) over time for different window types. Data were present as mean SEM. **e.** SHG (purple) and THG (cyan) imaging of skull from a mouse with OC window over 21 days. x-z images (top row) show the increase of skull thickness. The x-y image of osteocytes at the top of skull (indicated by top red dashed line) from THG signal channel is stable for 21 days (middle row). Bottom of the skull and dura (indicated by bottom red dashed line) at day 0 starts to show different patterns at day 7 to day 21 (bottom row).

Notably, we found that the sealing material of the thinned skull window is critical to its performance. When the window was sealed with silicone gel (Kwik-Cast, WPI)^29^ instead of UV gel (Kafuter 303), fluorescence signals decreased much more rapidly within a week (Fig. S2) compared with other windows. TPEFM imaging of the window and underlying tissue revealed large numbers of GFP+ cells, including a large population of myeloid cells (Fig. S2a), such as monocytes and macrophages in the dura^30,31^, a phenomenon not observed in windows sealed with other materials. The recruitment of these cells introduces strong scattering and degrades imaging quality^32^. A possible explanation is that sealing with silicone gel fails to protect the thinned skull from infection as effectively as other sealing materials.

For the OC window, we noticed that the clearing effect is limited when vessel structures are present in the spongy bone (Fig. S3), a condition commonly observed in relatively aged mice^19,33,34^. To recover the clearing effect, vessels in spongy bone are required to be removed by thinning the skull. Furthermore, OC window preparation is time-consuming, especially for aged mice, while thinning this outer layer prior to optical clearing markedly accelerates the clearing process by physically reducing the dense bone thickness for chemical treatment. After thinning, the skull thickness is approximately 30–50□µm, which preserves mechanical strength while facilitating optical clearing and ultimately improving imaging quality (Fig.1b). Given these advantages, shortened clearing time from over one hour to about 10 minutes and compatibility with a broader age range, we designated this optimized approach as the thinned-clearing (TC) window to distinguish it from the OC window in this study. We found that the TC window increased fluorescence signal intensity by approximately 1.7-fold compared with the thinned skull window (Fig. 1a, b). For a quantitative comparison of the four window types, their fluorescence intensities relative to the thinned-skull window are shown in Fig. S4.

Having established a standard for signal intensity comparison, we next investigated the longevity of each transcranial window by longitudinally imaging the same brain sites. The pattern of cerebral vessels served as landmarks to locate the same imaging positions on different days. We observed a rapid decrease in GFP signal (Fig. 1c), particularly from fine microglial structures in the deep brain (Fig. S5), which coincided with a marked increase in the thickness of the second harmonic generation (SHG) signal at the inner skull surface (Fig. 1d, Fig. S6a-b, Fig. S7a-b). To confirm that the newly appearing SHG signal corresponds to regrown bone, we performed third harmonic generation (THG) imaging to visualize osteocyte structures using a three-photon microscope^35^. Representative results from an OC window are shown on Fig. 1e, as skull in OC window is thick for clearly display osteocytes. Consistent with our SHG findings (Fig. S6a-b and S7a-b), we observed a gradual increase in THG signal thickness at the inner skull layer (Fig. 1e, top row). Importantly, THG imaging clearly revealed mature osteocytes within the skull (Fig. 1e, second row) and clearly identified newly formed osteocytes located beneath original skull, coinciding with the newly appeared SHG signal region (Fig. 1e, bottom row). These observations demonstrate that new skull structures originate at the base of the original bone and contribute to the fluorescence signal decay observed with the OC window. Similar patterns of skull regrowth and GFP signal reduction were also detected across the thinned skull window, PoRTS, and TC window (Fig. S6 and Fig. S7), indicating that degradation of optical transmission in all transcranial windows are due to skull regrowth.

An intuitive approach to recover signal after skull regrowth in thinned skull windows is to re-thin the skull, as reported previously^15^. In this study, we quantitatively evaluated the effects of re-thinning. Mice with thinned skull windows were imaged longitudinally until day□14, when skull regrowth occurred and signal levels declined. After re-thinning, we observed only modest signal recovery compared to pre-procedure levels at the same imaging site (Fig. S8a-c). Importantly, the recovered signal did not reach the original baseline intensity, even though the skull was re-thinned to a level below its initial thickness (Fig. S8b-c), which also resulted in markedly weakened mechanical integrity. It is known that skull regrowth progresses from woven bone in the early stage to laminar bone at later stages, two structures with distinct optical properties^36,37^. A plausible explanation is that the newly regrown woven bone has inferior optical properties compared with mature laminar bone, thereby limiting signal recovery after re-thinning. For OC window, previous studies have suggested that re-clearing can restore transparency and recover signal intensity^38^. To test whether re-clearing could extend the longevity of the OC window, we repeated the clearing procedure after two weeks, when the signal had already decayed substantially. However, we observed little to no recovery of signal intensity following re-clearing (Fig. S9), indicating that the newly formed skull beneath is resistant to clearing and that extending OC window performance through re-clearing is unlikely.

### Optical properties of transcranial windows: aberrations and resolution

High-resolution brain imaging is essential for investigating fundamental biological processes and disease mechanisms in living mouse models^39,40^. However, all transcranial windows introduce aberrations and scattering that distort the excitation light wavefront, thereby limiting both imaging resolution and depth. Adaptive optics (AO) has recently been applied to microscopy to correct sample-induced aberrations and scattering, improving both resolution and depth of brain imaging^4,5,21,22^. In this study, we comprehensively evaluated high-resolution imaging performance and optical aberrations across thinned skull, PoRTS, OC, and TC windows using a state-of-art two-photon ALPHA-FSS system^21,22^ and a wavefront sensor-based AO-TPEFM^4^. We first performed high-resolution deep imaging through the different transcranial windows by correcting optical aberrations with the two-photon ALPHA-FSS system. Using the TC window as an example, with a skull thickness of approximately 20□µm, microglial processes became blurred at ∼200□µm below the pia and were unresolvable beyond 300□µm when using standard TPEFM (Fig. 2a-b). By applying direct focus sensing with the GFP fluorescence signal, two-photon ALPHA-FSS effectively compensated for sample-induced aberrations, leading to a marked increase in the fluorescence intensity of microglial processes (Fig. 2a-b). This enabled clear visualization of the finest microglial structures beyond 440□µm (Fig. 2b), demonstrating the capability of *in vivo* high-resolution imaging of the mouse cortex through the TC window across large volumes. Comparable improvements in resolution and imaging depth were also achieved in the thinned skull, PoRTS, and OC windows using two-photon ALPHA-FSS TPEFM (Fig. S10–S12), indicating the aberration correction capability of ALPHA-FSS in various transcranial windows for high resolution deep brain study.

**Figure 2:**
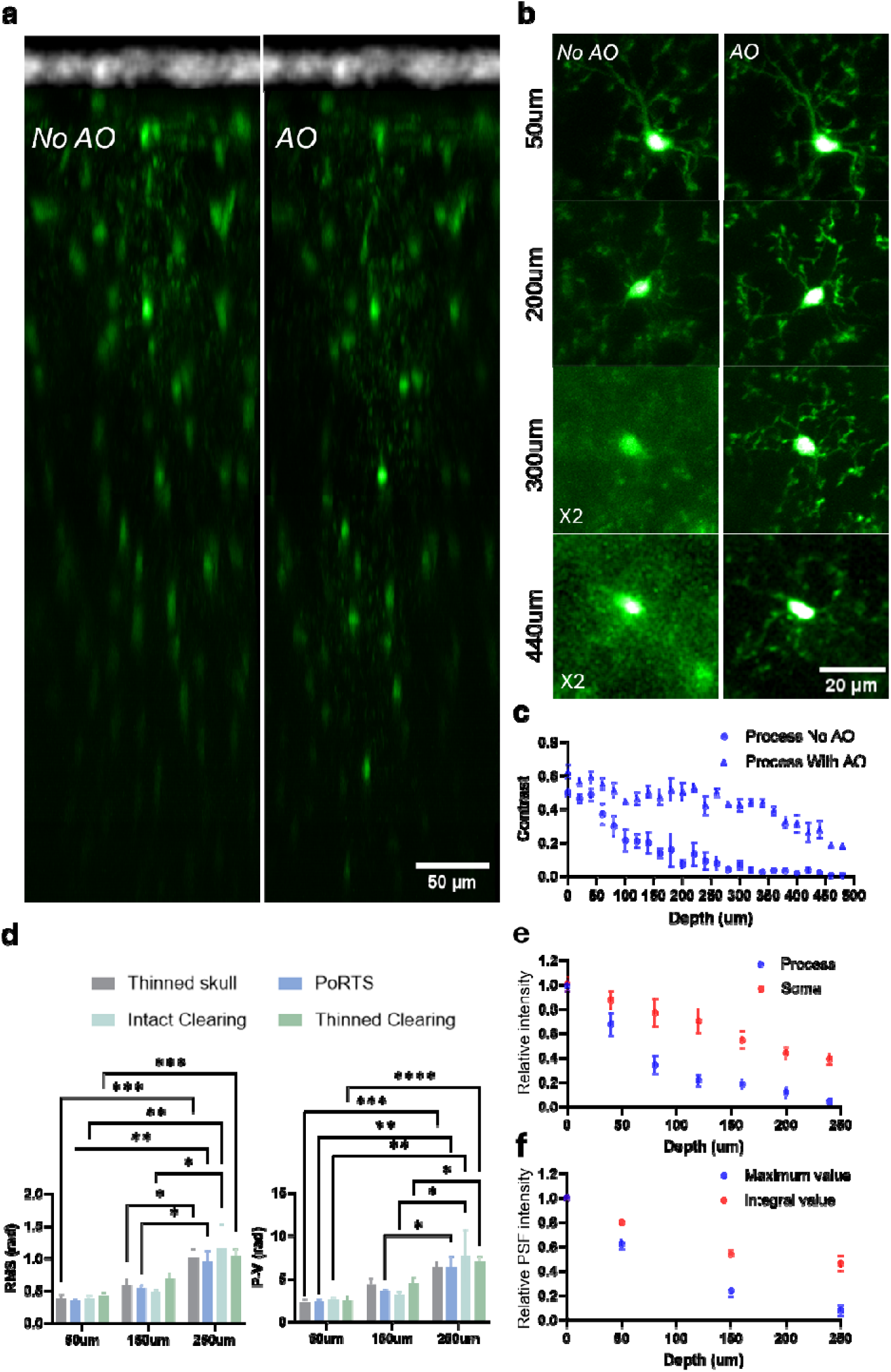
Imaging resolutions and optical aberrations of various transcranial windows. **a.** Two-photon image of TC window along y-axis on a *Cx3cr1^GFP/+^* mouse under system aberration correction and full AO correction by two-photon ALPHA-FSS system. **b.** Representative images of microglia at different imaging depths before and after AO correction. All the images are on the same gray scale. **c.** Contrast of microglia processes decay curve with imaging depth. Blue circle indicates the contrast decay curve without AO and blue triangle curve indicates the contrast decay curve with AO. Data were collected from one mouse each in the thinned-skull, PoRTS, OC, and TC groups, and are presented as mean□±□SEM. **d.** Comparison of aberration for different transcranial windows at different depths through a WFS based AO-TPEFM system (6 mice for thinned skull, 5 mice for PoRTS, 2 mice for TIS, 5 mice for TC). RMS value (left figure) and peak-valley value (right figure) are calculated through the measured wavefront. Data were present as mean ± SEM. Two-way ANOVA with Tukey’s multiple comparisons test. *p<0.05; **p<0.01; ***p<0.001; ****p<0.0001. **e.** Comparison of relative intensity of microglia processes and soma without two-photon ALPHA-FSS. Intensity values are normalized with the intensity of processes and soma from the same microglia image after two-photon ALPHA-FSS. **f.** Comparison of maximum intensity and integral intensity value of simulated PSF from the measured wavefront with WFS based AO-TPEFM. Intensity values are normalized with PSF intensity value from a perfect plane wavefront

Given the limited quantitative studies on TPEFM imaging resolution across different transcranial windows, we quantified resolution degradation with imaging depth through the thinned-skull, PoRTS, OC, and TC windows, using the high-resolution images produced by ALPHA-FSS as a reference standard. Here, fine microglial processes with diameters of approximately 1 μm were used as high-sensitivity structural indicators of resolution. As these processes became rapidly blurred with depth (Fig. 2b), it was difficult to assess resolution over 200 um depth by fitting the line profile of microglial processes as described previously^5,41^. Instead, we estimated resolution using the contrast of microglial processes (see Methods). Specifically, high-resolution reference images obtained with two-photon ALPHA-FSS were first used to generate accurate binary masks delineating microglial processes and background. These masks were then applied to images acquired with and without AO correction to calculate process contrast at multiple depths. The contrast degraded sharply within 200□µm, indicating that the effective high-resolution imaging depth without AO is less than 200□µm (Fig. 2c), consistent with visual assessment. With AO correction, however, process contrast was significantly enhanced and maintained down to 450□µm in depth (Fig. 2c).

To understand the rapid decrease in imaging resolution with increasing imaging depth, we studied optical aberrations across various transcranial windows using wavefront sensor–based AO TPEFM, which directly provides continuous wavefront measurements for analysis^4^. Here, GFP-labeled microglia at depths of 50□µm, 150□µm, and 250□µm were used as guide stars. Deeper measurements were excluded because the increased background noise beyond 250□µm substantially compromised the accuracy of wavefront sensing. Although a longer-wavelength fluorescent guide star could, in principle, extend the measurable depth, the present analysis using GFP fluorescence was restricted to within 250□µm^4^. We found that both peak-to-valley (P–V) and root-mean-square (RMS) wavefront error analyses, commonly used indicators of aberration, showed no significant differences among window types (Fig. 2d). However, aberrations increased markedly with imaging depth, consistent with the rapid deterioration of resolution in deeper regions (Fig. 2b,c). This is likely due to the larger skull area encompassed by the imaging laser beam at greater depths, which introduces additional optical aberrations. Representative data for the thinned-skull, PoRTS, OC, and TC windows are provided in the Supplementary Information (Fig. S13-S16).

It is worth noting that although imaging resolution declined rapidly within 200□µm, microglial somas remained clearly resolvable beyond 400□µm even without AO correction (Fig. 2b). To quantify this observation, we calculated the relative fluorescence intensities of somas and processes in non-AO images, normalized to AO-corrected references across depths (see Methods), and found that soma intensities decayed much more slowly than those of the processes (Fig. 2e). To understand why somas are less sensitive to aberrations^42^ than microglial processes, we reconstructed three-dimensional two-photon point-spread functions (PSFs) at varying depths using wavefront data from the wavefront sensor-based AO-TPEFM. The maximum intensity of the aberrated PSF and its integrated intensity within a soma-sized volume (15×15×15 *um*^3^) were normalized to those of an ideal PSF to simulate fluorescence changes in microglial processes and somas, respectively (Fig. 2e). We observed that integral value of aberrated PSF decays slower than the maximum value of aberrated PSF (Fig. 2f), consistent with the experiment results (Fig. 2e). These findings indicate that the integrated PSF value, corresponding to larger structures in tissue, is less sensitive to aberrations than its peak intensity, thereby explaining the slower intensity decay of microglial somas in deeper tissue.

As skull regrowth gradually attenuates fluorescence intensity, we further assessed the extent to which two-photon ALPHA-FSS TPEFM could correct aberrations introduced by newly formed bone and thereby extend the imaging duration. We performed longitudinal imaging of *Cx3cr1^GFP/+^*mice through a TC window at the same site over three weeks. At day 0, AO correction produced a marked improvement in signal, with both intensity and resolution of microglia enhanced at depths beyond 400□µm (Fig. 3a–d). By day□21, however, GFP fluorescence had substantially degraded due to skull regrowth (Fig. 3e). Under these conditions, accurate PSF measurement became difficult, thereby reducing the effective correction depth. Although AO still yielded some improvement in resolution and fluorescence intensity, both the correction efficiency and the maximum imaging depth were markedly diminished compared with day□0 (Fig. 3f–h). These findings indicate that the ability of ALPHA-FSS to compensate for skull-induced aberrations becomes increasingly limited as skull regrowth progresses.

**Figure 3.**
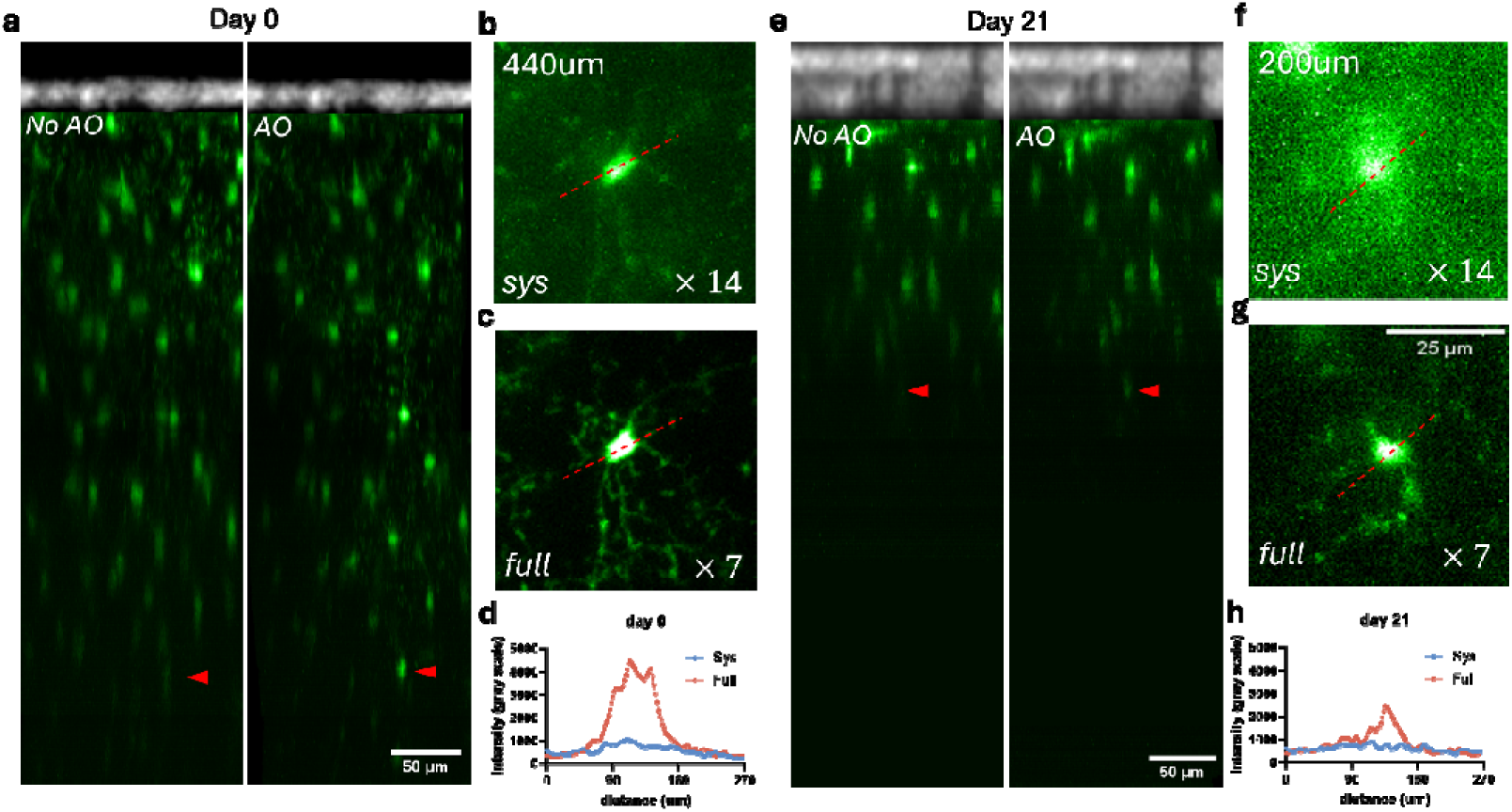
ALPHA-FSS TPEFM imaging during skull regrowth. **a.** Deep brain imaging at day 0 after thinned skull surgery without and with AO correction based on two-photon ALPHA-FSS. Same skull images are shown by the SHG signal (gray) and microglia is shown by GFP (green). **b-c**. Image of microglia at 440 µm below pia without and with AO correction. **d**. Comparison of line profile along the microglia soma with and without AO correction along the red dashed line in Fig.3b and 3c. **e.** Deep brain imaging at day 21 after thinned skull surgery without and with AO correction based on two-photon ALPHA-FSS. **f-g**. Image of microglia at 200 µm below pia without and with AO correction. **h**. Comparison of line profile along the microglia soma with and without AO correction along the red dashed line in Fig. 3f and 3g. Gray scale is the same for all the images.

### Long-term transcranial windows with skull regrowth suppression

As demonstrated by the results above, skull regrowth is the primary factor limiting window longevity. A direct and practical strategy to extend imaging duration is therefore to develop methods that suppress this regrowth. Glucocorticoids (GCs), a class of steroid hormones involved in diverse physiological processes such as metabolism, mood regulation, and development^43,44^, are well documented to cause osteoporosis and growth retardation^23,25^. Their growth-suppressing effects act on all bone cell types, promoting osteoclast activity and bone resorption while inhibiting osteoblast-mediated matrix synthesis^23,24,45^. To test whether GC treatment could provide a simple means of inhibiting skull regrowth and stabilizing its optical properties, we first applied an off-the-shelf commercial synthetic GC ointment to the thinned skull immediately after surgery (see Methods). After preparing the thinned skull window and finishing *in vivo* imaging on day 0, a droplet of dexamethasone ointment was placed directly on the exposed skull and sealed the ointment with removable UV gel. For subsequent imaging sessions, the UV gel and residual ointment were removed before performing *in vivo* imaging with TPEFM. Remarkably, dexamethasone treatment effectively inhibited skull regrowth for over four weeks (Fig. 4a,b; Fig. S17). We also tested an endogenous GC derivative, hydrocortisone ointment (see Methods), and observed a similar inhibition of skull regrowth (Fig. S18). Correspondingly, GFP signal intensity remained stable as long as skull thickness was preserved, in contrast to the rapid degradation of the pure thinned-skull window within one week (Fig. 4a,c). These findings demonstrate that local applications of GC ointment can effectively suppress skull regrowth and thereby extend the imaging duration achievable with the thinned skull window. It should be noted that the PoRTS window requires cyanoacrylate cement to reinforce the enlarged thinned skull^16^, whereas the OC and TC windows require a specialized UV-curable adhesive (S3) to retain the clearing agents within the skull^17^. Because these sealing methods preclude local GC delivery, only the thinned-skull window is suitable for GC treatment to suppress skull regrowth.

**Figure 4.**
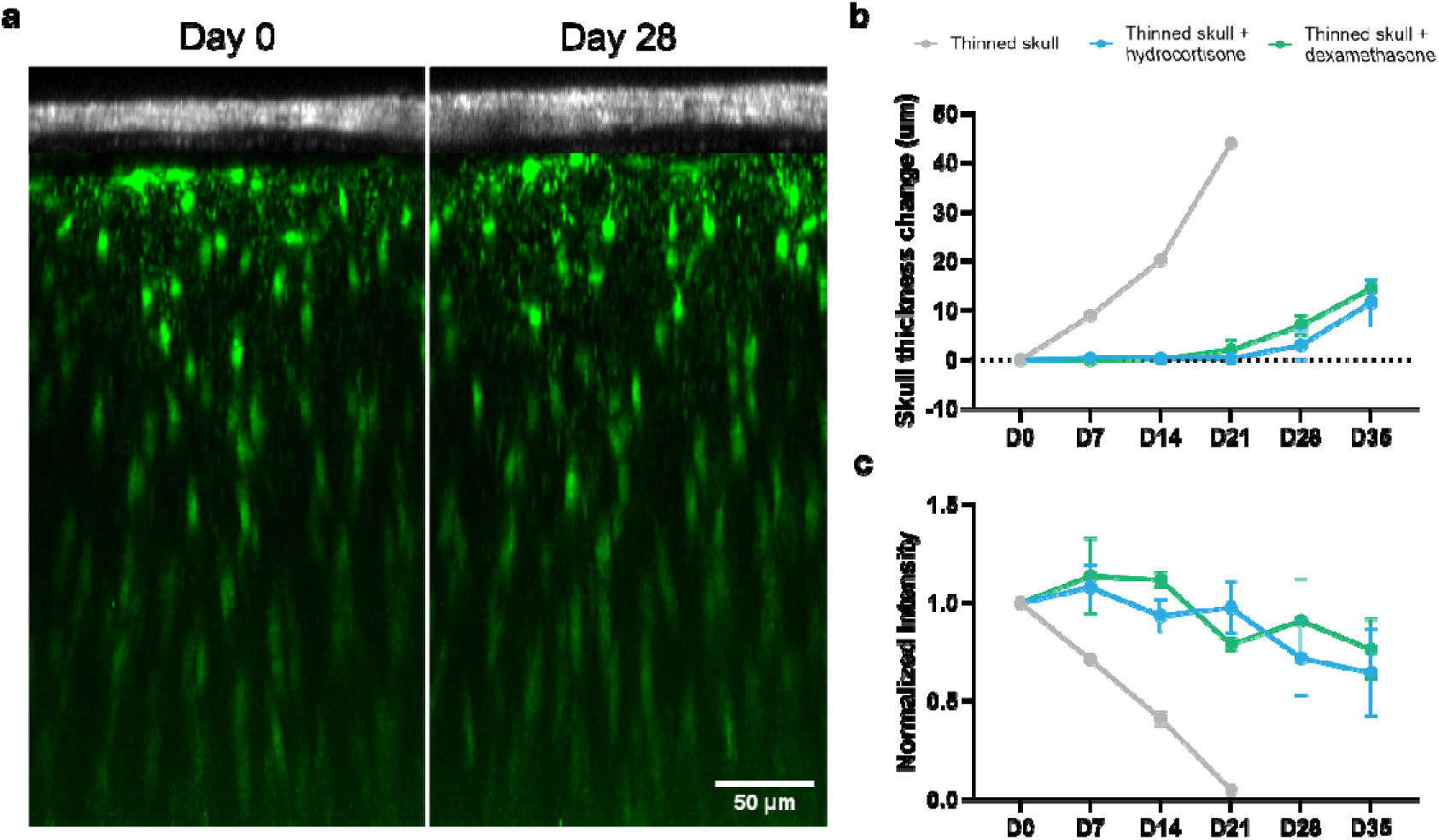
Inhibition of skull regrowth by local glucocorticoid application. **a**. Long-term imaging of thinned skull window with locally applied off-the-shelf dexamethasone ointment over 28 days. Skull is shown by the SHG signal (gray) and microglia is shown by GFP (green). **b-c.** Statistics of skull thickness changes at different days (upper) and statistics of normalized GFP fluorescence intensity at different days (lower) for thinned skull window with hydrocortisone and dexamethasone ointment. Data were present as mean ± SEM.

While skull regrowth was effectively inhibited by off-the-shelf GC ointment, we observed displacement of microglia position (Fig. 5a-b) and morphological alterations in microglia (Fig. 5c-e) within the superficial cortex (<60□µm) following application of off-the-shelf dexamethasone or hydrocortisone ointment to the thinned skull. We then evaluated changes in microglial morphology after GC treatment using two quantitative indicators: ramification index and surveillance area (see Methods). We found that microglia exhibited increased ramification indices (Fig. 5c) and surveillance areas (Fig. 5d) after application of dexamethasone ointment, an effect that was not observed in the hydrocortisone ointment group. To assess the displacement of microglia, we classified microglia that persisted in the same position from day□0 to day□7 as “stationary” and those that disappeared or newly appeared as “dynamic”. This analysis revealed fewer stationary and more dynamic microglia after drug application, whereas most microglia in controls remained stationary, consistent with previous studies (Fig. 5e)^46^. Moreover, in a subset of mice we occasionally observed stronger microglial activation, as indicated by a marked decrease in ramification index after 7□days of local dexamethasone or hydrocortisone application (Fig. S19), which may be attributable to differences in GC permeability across thinned skulls. These activation effects on microglia are likely due to the small molecular size of glucocorticoids, which enables them to penetrate the meninges, diffuse into the brain parenchyma, and perturb microglial physiology^47,48^.

**Figure 5:**
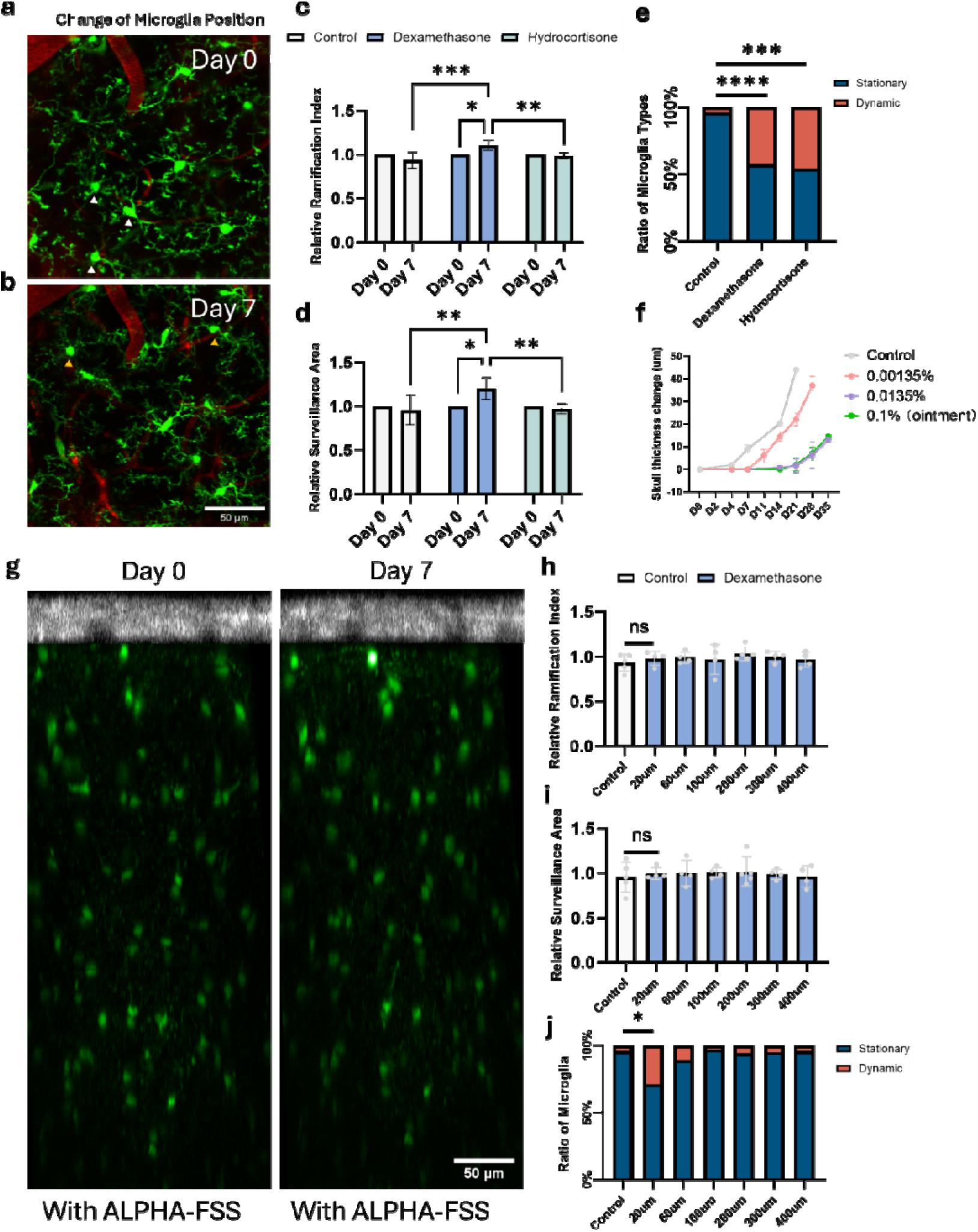
Glucocorticoid effects on microglia. **a-b.** Microglia changes its position before (a) and after 7 days (b) of delivery of dexamethasone or hydrocortisone ointment. White arrow indicates the loss of microglia and orange arrow indicates the newly appeared microglia. **c.** Relative ramification index changes of same microglia after 7 days for control group and hydrocortisone/dexamethasone ointment delivery group. Ramification index of each microglia is normalized to its ramification index of day 0. **d.** Relative surveillance area of same microglia after 7 days for control group and hydrocortisone/dexamethasone ointment delivery group. Surveillance area of each microglia is normalized to its surveillance area of day 0. **e.** Contingency table of ratio between stationary microglia and dynamic microglia after 7 days for control group and hydrocortisone/dexamethasone ointment delivery group. **f.** Skull thickness changes of different dexamethasone concentration. **g.** High-resolution deep brain imaging of Cx3Cr1-GFP mouse with 0.0135% dexamethasone delivery for 7 days using two-photon ALPHA-FSS. Change of microglia is analyzed based on high resolution imaging on day 7 (right) compared with day 0 (left). **h.** Comparison of relative ramification index changes of same microglia between control group and 0.0135% dexamethasone group at different depths. **i.** Comparison of relative surveillance area changes of same microglia between control group and 0.0135% dexamethasone group at different depths. Data were present as mean ± SD. RM Two-way ANOVA multiple comparisons test. Only the first ns is shown. **j.** Comparison of ratio of stationary and dynamic microglia changes between control group and 0.0135% dexamethasone group at different depths. Fisher’s exact test is used. *p<0.05; **p<0.01; ***p<0.001; ****p<0.0001. ns is not shown.

To reduce GC effects on microglia, we examined its concentration dependence to determine whether skull regrowth could be inhibited at lower GC concentrations without adversely affecting microglia. First, a small piece of medical sponge was used as a reservoir to deliver dexamethasone solution to the thinned skull at various concentrations. Briefly, the dexamethasone-loaded sponge was placed on the exposed thinned skull area, and removable UV gel was then applied on top of the sponge to seal the window, following the same procedure used with the ointment (Fig. S20). *In vivo* imaging was performed weekly; during each session, the UV gel and sponge were removed and then reapplied afterward. We found that when the dexamethasone concentration was reduced to 0.00135%, its inhibitory effect on skull regrowth began to diminish. By contrast, a moderate concentration (0.0135%), approximately 10-fold lower than that of the off-the-shelf commercial dexamethasone ointment (0.1%), successfully inhibited regrowth for up to 28 days (Fig. 5f, Fig. S21). *In vivo* ALPHA-FSS TPEFM imaging, which enabled analysis of both microglial morphology and displacement at depths of up to 400□µm below the pia (Fig. 5g), further revealed that, at this moderate concentration, dexamethasone no longer altered microglial morphology in the superficial cortex, and microglia in deeper brain regions remained unaffected (Fig. 5h,i). While the proportion of stationary microglia in the superficial cortex remained above 70% (Fig. 5h), no statistically significant changes in microglial position were observed at depths below 60□µm (Fig. 5j). Together, these findings indicate that the effects of dexamethasone on microglia are dose-dependent, and that carefully controlled lower concentrations can minimize its impact on microglia. However, we were unable to identify a dexamethasone concentration that both effectively inhibited skull regrowth and completely eliminated effects on microglia. Future studies should therefore focus on identifying pharmacological agents that suppress skull regrowth while exerting minimal influence on brain physiology.

Having demonstrated that GCs can inhibit skull regrowth and thereby extend the imaging duration achievable with the thinned-skull window, we next examined the feasibility of establishing a chronic transcranial window with high optical transmission for longitudinal *in vivo* brain imaging. In recent years, numerous studies have highlighted the potential of hydrogels as biocompatible materials for drug delivery. In particular, many commercially available hydrogels exhibit excellent optical properties and can readily load and release drugs^49–51^, making them promising candidates for use in chronic transcranial imaging windows. In our study, we replaced the medical sponge with a hydrogel loaded with dexamethasone at the moderate concentration (0.0135%). Specifically, we employed a commercial UV-photocrosslinked methacrylated hydrogel composed of hyaluronic acid methacrylate (HAMA) and the photoinitiator lithium phenyl-2,4,6-trimethylbenzoylphosphinate (LAP), which was loaded with dexamethasone solution and applied to the surface of the thinned skull. To ensure stability and prevent dehydration, we designed a custom head mount in which a round coverslip of appropriate size was placed on top of the hydrogel and secured to the mount with removable UV gel, thereby preventing water evaporation from the hydrogel (Fig. 6a, Fig. S22).

**Figure 6:**
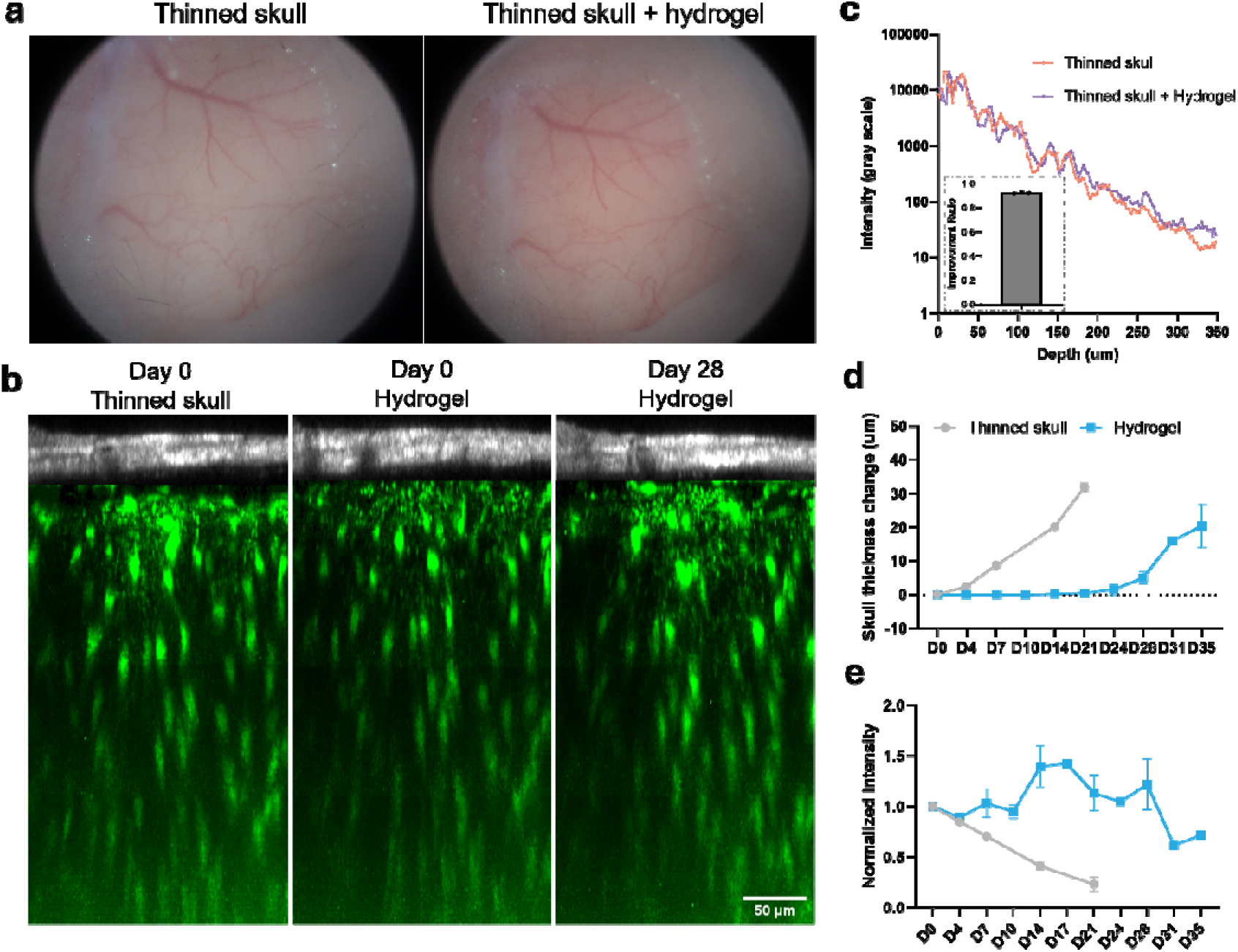
Hydrogel-based chronic transcranial optical window for skull regrowth inhibition. **a.** Wide field image of thinned skull window and thinned skull window sealed with hydrogel. **b.** Maximum projection image of thinned skull window (left), thinned skull + hydrogel window (middle) and 28 days after the thinned skull surgery (right) along y-axis on a Cx3Cr1-GFP mouse. Skull is shown by the SHG signal (gray) and microglia is shown by GFP (green). **c.** GFP intensity changes for a same mouse before and after hydrogel window is applied. Statistics of relative intensity changes after hydrogel window is applied in the sub-figure. **d.** Statistics of skull thickness changes at different days between thinned skull window and hydrogel window. **e.** Statistics of relative intensity changes at different days between thinned skull window and hydrogel window. Data were present as mean SEM on 4 mice.

We first compared imaging quality before and after applying the hydrogel window and observed negligible signal loss (Fig. 6b, c), indicating that the HAMA hydrogel maintained high transparency during two-photon imaging. We next evaluated the efficacy of the dexamethasone-loaded hydrogel in inhibiting skull regrowth by longitudinally tracking skull thickness at the same site. Notably, the HAMA hydrogel exhibited shrinkage over 4 days due to dehydration^52^, resulting in a loss of tight contact with both the coverslip and the skull. This occasionally led to the formation of air bubbles within the window, which interfered with imaging. To address this issue, we refreshed the window every 3 or 4 days to maintain imaging quality and ensure continuous contact between the hydrogel and the skull for effective drug release. Remarkably, skull thickening was suppressed for over 28 days without appreciable signal loss, demonstrating the effectiveness of the hydrogel in delivering dexamethasone to inhibit regrowth (Fig. 6d, e, Fig. S23). However, skull regrowth resumed and signal level decreases after 28 days (Fig. 6d-e), similar to the results obtained with the sponge loaded with dexamethasone solution (Fig. 5f). In future work, new hydrogel-based materials or alternative approaches with reduced dehydration could be explored to extend the duration of the optical74 imaging window^53^.

## Discussion

Optical windows for transcranial brain imaging are a critical technology for investigating brain physiology and disease in mouse models. With recent advancements in multiphoton microscopy, imaging depth has markedly improved across several types of transcranial windows, including the thinned skull, PoRTS, and OC windows. Two fundamental questions commonly considered by researchers are how long these windows remain effective and how deep high-resolution images can be obtained. In this study, we addressed these issues by quantitatively evaluating both the imaging duration of different transcranial windows and the achievable imaging depth using advanced adaptive optics two-photon microscopy. It is important to emphasize that imaging quality is influenced by multiple factors, including skull thickness, the age of the mice, and the precision of thinned-skull surgery. Taking together, the study yielded the key insights into the performance of different transcranial windows. Furthermore, we explored an approach to extend imaging duration, maintaining stable fluorescence intensity for up to four weeks, by inhibiting skull regrowth through local delivery of glucocorticoids. This work paves the way for future efforts to further optimize chronic transcranial optical windows for *in vivo* mouse brain imaging.

As the earliest form of transcranial window, the thinned-skull window offers advantages of simplicity and reduced invasiveness compared with open-skull surgery. Although skull regrowth has been reported to cause signal degradation in the thinned-skull window^15^, we identified an additional limiting factor, dural inflammation, that substantially shortens window duration, restricting imaging to less than one week. Notably, dural inflammation frequently arises when the thinned-skull window is sealed with silicone gel (Kwik-Sil), as reported in previous studies^29^, whereas this complication can be prevented when the window is sealed with UV gel. A plausible explanation is that infection from the external environment promotes cell recruitment and inflammation, suggesting that the barrier and protective properties of silicone gel are less effective than those of UV gel. This observation also explains why no dural inflammation was detected in other transcranial window preparations, all of which employed stronger protective sealants such as cyanoacrylate cement (PoRTS) or UV gel (OC, TC).

Our study revealed that all transcranial windows, including thinned skull, PoRTS, OC, and TC, undergo skull regrowth, which represents the fundamental mechanism limiting their long-term performance. Although previous work suggested that PoRTS could enable extended imaging by inhibiting bone regrowth through fusion of the window with cyanoacrylate cement^16^, the discrepancy with our findings may arise from methodological differences in assessing signal and image quality as well as in experimental design. Specifically, the previous study measured skull thickness for up to 7 days in two mice^16^, whereas our analysis involved longitudinal continuous measurements of stable GFP signal sources over four weeks. We found that skull regrowth is relatively modest during the first week following thinned-skull surgery or the OC procedure; however, it accelerates in subsequent weeks, leading to a rapid decline in signal intensity and image quality. We found that skull regrowth originates from the inner surface of the remaining bone. This occurs because the periosteum of the superficial compact bone, which is essential for regeneration and remodelling of bone^54^, is removed during window preparation, whereas the periosteum on the inner surface of the skull remains intact^55^. Consequently, the PoRTS window should be used with caution for imaging procedures extending beyond one week, particularly in experiments requiring high resolution and depth. Notably, we also observed skull regrowth in the OC window despite the skull remaining intact. Furthermore, we demonstrated that the newly formed bone induces signal degradation that cannot be reversed by re-clearing.

It is well established that exposure to high doses of glucocorticoids (GCs) induces osteoporosis in various bone tissues^23,24^. In this work, we leveraged this side effect on bone growth to study whether GCs could be used to inhibit skull regrowth and thereby elongated transcranial window performance. We first demonstrated that off-the-shelf glucocorticoid ointments effectively suppressed bone regrowth, leading to stable microglial GFP fluorescence signals over several weeks. However, given the small molecular weight of GCs, they can penetrate the blood-brain barrier and diffuse into the brain, thereby influencing superficial microglia^45^. A known drawback of GCs is their dose-dependent effect on microglia in the superficial cortex^56^. By administering an appropriately low concentration of dexamethasone, we were able to prevent morphological alterations in microglia, although slight displacement was still observed, indicating minimum drug effects. Importantly, an unchanged morphological index does not necessarily indicate that microglia is unaffected; additional biochemical or transcriptomic analyses will be required to more comprehensively assess the drug’s impact on microglia and other brain cell types in the future work. Finally, we found that skull regrowth becomes apparent about four weeks after GC treatment, likely due to GC resistance arising from long-term exposure and activation of the glucocorticoid receptor (GR)^57,58^. Nonetheless, our findings underscore the importance of suppressing skull regrowth to achieve long-term imaging with transcranial windows. Therefore, a key future research direction is the identification of pharmaceutical compounds that specifically inhibit bone regrowth while remaining confined within the skull, without affecting the meninges and brain.

One of the most promising results from this study is that hydrogels loaded with dexamethasone can serve as effective sealing agents, simultaneously inhibiting skull regrowth and maintaining optical transparency for high-quality imaging. The primary limitation of currently available commercial hydrogels is shrinkage caused by dehydration^52^, which necessitates window replacement or refreshing every 3-4 days. Currently, numerous studies have been focused on developing anti-dehydration hydrogels that retain optical transparency and exhibit extended lifespans^53^. Future work will focus on exploring such hydrogels with effective drug delivery, ensuring they preserve the brain’s physiological environment while also enabling longer intervals between window refreshment.

## Methods

### Animal

All the animal experiments were conducted on adult *Cx3cr1^GFP/+(^B6.129P2(Cg)-Cx3cr1tm1Litt/J)* mice. Young mice are around 2-months old, and aged mice are around 5-months old. All animal procedures were conducted in accordance with the guidelines of the Animal Care Facility of the Hong Kong University of Science and Technology (HKUST) and received approval from the Animal Ethics Committee at HKUST.

### Thinned skull surgery

To conduct thinned skull surgery, a general procedure was conducted to establish a head-fixed surgery condition. The mice were anesthetized by intraperitoneal injection Ketamine–xylazine mixture (KX) solution. The thinned-skull preparation was slightly modified from a previous protocol^15^. After removing the scalp and periosteum around the target region (around 10 mm diameter), the tissue adhesive (3M Vetbond) was applied to seal the edge between remained scalp and skull. Then, metabond (C&B Metabond, Parkell) was applied on the skull except the target imaging region. A custom-made ring holder was sticked on the metabond layer by cyanoacrylate cement. The gap between holder and metabond was then sealed by filling dental cement.

To thin the skull, a 500 μm carbon steel burr attached to a rotatory high-speed drill was used to gently thin a circular region (1-1.5 mm in diameter) with the center at stereotactic coordinate (3 mm, 3 mm) laterally and posterior to the bregma point with saline filled the pool enclosed by the holder. After removing the majority of the middle spongy bone, a microsurgical blade (No. 6961, Surgistar) was used to carefully thin the skull further to about 20–40 μm. Surface irregularities were reduced by occasionally changing the thinning direction of the surgical blade. Finally, a biocompatible sealant mixture (Kwik-Cast, WPI), which can be peeled off before the imaging experiment, was applied to cover the thinned-skull window.

### PoRTS surgery

The PoRTS surgery preparation was conducted using previously described method^16^. Following the thinned-skull surgery, the thinned area was cleaned with saline solution to maintain moisture. The skull was then sequentially polished with slurry of tin oxide powder. A tiny drop of cyanoacrylate cement was placed on the skull surface and then immediately covered with a 4-mm circular coverslip. It was essential to quickly adjust the coverslip before the cement dried, ensuring proper positioning. Careful control of the cement drop was necessary to minimize the thickness of the window, as a thick cement-glass interface could lead to significant optical aberrations. During imaging, the correction ring can be adjusted to a proper position, guided by fluorescent signal intensity, thereby minimizing any aspherical aberrations.

### Optical clearing window & Thinned-clearing window

The Optical clearing window preparation was performed as previously described^17^. All the chemicals required for clearing process(S1, S2, and S3) were bought from Jarvis (Wuhan) Biopharmaceutical Co., Ltd, China. The optical clearing window was established on an intact skull following the head-fixed surgery, while the thinned-clearing window requiring optical clearing after the thinned-skull surgery.

To perform the optical clearing, S1 solution was applied on the dry skull and filled the pool enclosed by the holder. A fine-tip cotton swab was used to gently massage the skull. The S1 solution was replaced every 3 minutes until the skull surface in the target region transitioned from white to semi-transparent, allowing for clear observation of the underlying vessels. The clearing time is highly dependent on skull thickness, typically within 10 minutes for skull thicknesses less than 50 µm, but it can exceed half an hour for thicker skulls such as an intact skull. After achieving the desired clarity, the S1 solution was wiped away, and S2 was applied to the skull. The solution was allowed to sit for approximately 2 minutes until the tiny “bubble-like” structures in the skull disappeared, as observed through a stereomicroscope. Subsequently, most of the S2 was wiped off, and S3 was immediately applied to the skull. It is important to avoid completely removing S2 or exposing the skull to air, as this may cause the skull to become opaque again. If whitening occurs, the clearing procedures should be repeated. Use tweezers to remove the bubbles. The thinned region was covered with a 4-mm circular coverslip. Cotton was used to absorb excess liquid, allowing the coverslip to rest closely against the skull to minimize window thickness, as a thick S3-glass interface could lead to significant optical aberrations. Finally, used 405 nm LED to solidify S3 for 2 mins. Intermittently switched the LED every second to ensure uniform curing of S3.

### Administration of hydrocortisone and dexamethasone

Off-the-shelf dexamethasone (0.1% w/w dexamethasone, sterile ophthalmic ointment, TOBRADEX) and off-the-shelf cortisol (1% w/w hydrocortisone, TaiGuk HC) were used by applying a droplet of ointment on the exposed thinned skull. Gently smear the ointment so that all the skull area is covered. Then, UV glue (Kafuter 303) was applied on the ointment to seal the window and solidified using 405 nm LED for 20-30s.

For locally treatment of mice with dexamethasone solution of different concentration, the dexamethasone was dissolved in saline. The concentration was based on typical plasma levels from global treatment (1-10 mg/kg/day)^25^. For a 25 g mouse, the dexamethasone concentration should be between 0.0125-0.125 mg/ml. A piece of hemostatic sponge was soaked with dexamethasone solution and covered all the exposed skull. Then, to seal the window, UV glue (Kafuter 303) was applied on the sponge and totally sealed the window. Then the UV glue was solidified using 405 nm LED for 20-30s. The UV glue cover can be easily removed by tweezers before *in vivo* imaging.

### Optical window with dexamethasone loaded hydrogel

HAMA (50 kDa) and LAP were obtained from YuXi Biology, China. The hydrogel was prepared following the protocol provided by YuXi Biology. First, 0.05 g of LAP was dissolved in 20 ml of PBS in a 40°C water bath for 15 minutes. Then, 40 mg of water-soluble dexamethasone (Sigma-Aldrich) was added to the LAP solution. Next, HAMA was added to the LAP solution at a concentration of 4% (w/v) and stirred for 30 minutes. Approximately 40 µl of the hydrogel solution was then dropped onto the skull, and a 10 mm coverslip was carefully placed on top. The hydrogel was solidified using a 405 nm LED for 20 seconds. The edges of the coverslip were sealed with UV glue (Kafuter 303) and cured with the 405 nm LED for 20-30 seconds.

### *In vivo* imaging with TPEFM

After being anesthetized with ketamine and xylazine, mice were mounted on an angle adjuster (MAG-2, NARISHIGE, Japan). Texas Red (70KDa-Dextran, 50 µl, 2mg/100ul) was injected retro-orbitally into the mouse to label the vessel structures, which will be used as the reference for longitudinal imaging at the same site. 920nm femtosecond laser (FF Ultra 920, TOPTICA) was focused in the brain using a high numerical aperture (NA) objective (XLPLN25XWMP2, Olympus). The objective was mounted on a motorized linear actuator (LNR50SEK1, Thorlabs) for axial sectioning. The pupil of objective was underfilled to achieve an effective NA of 0.7. Imaging power was empirically set at different imaging depth: 10 mW (0-50 um), 20 mW (50-150 um), 40 mW (150-250 um) and 80 mW (250-350 um, with the same power configuration used for all mice.

The emitted fluorescence signals were separated by a dichroic mirror (Semrock, FF560-Di01-25x36). Suitable band-pass and short-pass filters were placed in front of two photomultiplier tubes (Hamamatsu, H7422-40) to selectively detect the SHG/Texas Red and GFP signals. The current signal from the PMT was converted to a voltage signal using a transimpedance current amplifier (DHCPA-100, Femto; SR570, Stanford Research).

The fluorescence signal intensity of each frame was calculated as the average fluorescent signal intensity of the brightest 0.3% pixel. We calculated the fluorescence signal intensity of each frame and plot as the function of the depth. After that, the power changes at different depths is then normalized to reach a continuous signal decay curve. We expressed the logarithm of fluorescence signal intensity *I* as a function of depth *d*: *I*=*α* · *d* + *I*_0_, where *α* indicates how strong the signal decay as the depth *d* increase, and *I*_0_ is the intercept which represents the globe fluorescence signal intensity.

### Aberration measurement with wavefront sensor based AO-TPEFM

A wavefront sensor based AO-TPEFM system^4,59,60^ was used to measure the aberration of different transcranial windows as it can output continuous wavefront for easier analysis peak-valley (P-V) value and root-mean-square (RMS) value of wavefront. Briefly, the emitted fluorescence in brain followed the reverse path of the excitation laser and was descanned by the galvo mirrors and then reflected by the deformable mirror (DM, Alpao, DM97-15). It will be separated by from the femtosecond laser by a dichroic mirror (Semrock, FF705-Di01-25*36) and then relay to the microlens array (SUSS MicroOptics, 18-00197) of a customized Shack-Hartman wavefront sensor (SHWS). The microlens array focuses the fluorescence onto an electron-multiplying charge-coupled device (Andor iXon3 888), and the shifts of the focus spots are used to calculate the wavefront. We measured both P-V and RMS values of measured wavefront from thinned skull, PoRTS, TIS, and TC windows at depths of 50 μm, 150 μm, and 250 μm. Aberration measurements from the guiding star in deeper brain regions were not taken, as scattering significantly affects the accuracy of the wavefront sensor^4,5^.

### *In vivo* imaging with ALPHA-FSS Two-photon microscope system

A customized ALPHA-FSS two-photon system was used to robustly compensate aberrations and scattering and achieve high resolution deep brain imaging. The principle of ALPHA-FSS has been discussed in previous work^5,22^. Briefly, our two-photon ALPHA-FSS system consists of three key modules: direct focus sensing, conjugate AO and remote focusing models. A 920 nm femtosecond laser (Coherent, Axon 920) is used for two-photon excitation. A spatial light modulator (SLM) is placed in the conjugation plane of the skull to achieve conjugate AO configuration. To measure the scattered and aberrated E-field PSF, a weak scanning beam which raster scanned over a small field of view (FOV) is introduced to interfere with a strong stationary beam. The stationary beam is parked on a fluorescence guiding star which is the soma of microglia in this work. Thus, the two-photon (2P) excited fluorescence signal at scanning coordinate x is given by:

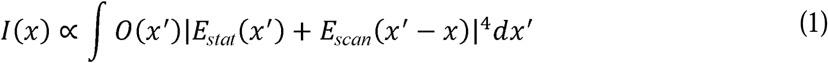

Where *E_stat_* and *E_scan_* is the complex-valued E-field PSF of the stationary and scanning beam and *O*(*x*’) is the real-valued object function related to the fluorophore’s distribution in the focal plane. Using an acoustic optical modulator (AOM, AOMO 3080-122, Crystal Technology) driven by a radio frequency (RF) signal generator (AODS20160-6 8, Crystal Technology) with frequency *ω*, the fluorescence signal of the interference beams is oscillating at frequency *ω*. For simplicity, we denote the complex amplitude of E-field PSF as *E_stat_* (*x*’) and *b* = *E_scan_* (*x*’ – *x*) and set the ratio between the two beams such that the stationary beam is much stronger than the scanning beam, 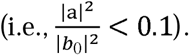 Then oscillating part at frequency *ω* in equation (1) can be approximated as:

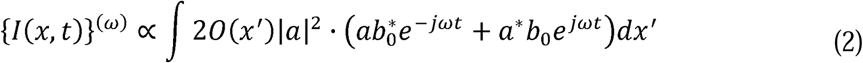

Using a lock-in amplifier with a reference signal at frequency *ω*, the point spread function can be demodulated as:

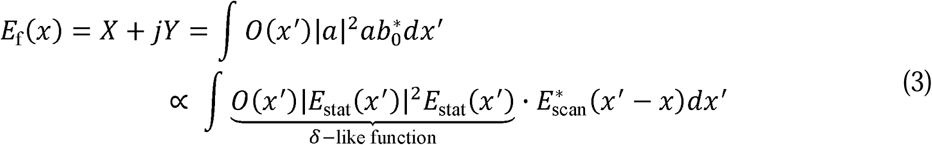

Where *X* and *Y* are in-phase and quadrature parts output of the lock-in amplifier. Therefore, the correction wavefront can be calculated by using the two-dimensional Fourier transform of the measured E-field PSF and applied to the SLM. An electrical tunable lens (ETL), placed in the conjugation plane of the pupil, is used to adjust the focal plane in brain without disrupting the conjugation relationship between SLM and skull.

Texas Red (70KDa-Dextran, 50 µl, 2mg/100 µl) was injected retro-orbitally into the mouse to label the vessel structures, which will be used as the reference for longitudinal imaging at the same site. For high resolution imaging with two-photon ALPHA-FSS, three times of PSF measurements are typically chosen at the depths of 150 – 250 µm, 250 – 350 µm and over 350 µm to cover the whole imaging volume. The excitation power starts from 30mW to maximum 150mW depends on the GFP signal intensity at different depth. The power configuration at different depths is recorded on day 0 and the same power configuration is used on different days for each mouse.

### Third harmonic generation microscope

For third harmonic generation (THG) imaging, the mice were anesthetized via intraperitoneal (i.p.) injection of a ketamine–xylazine mixture and then mounted on a head-holding stage with angle adjusters. The mouse skull was aligned to be perpendicular to the objective axis by using THG signal of skull as guidance. For longitudinal imaging, the same region was relocated using THG signal of blood vessels as guidance. Finally, volumetric imaging was performed from the skull surface to the superficial brain.

The excitation source is a two-stage optical parametric amplifier (Orpheus-F, Light Conversion) pumped by a 40 W femtosecond laser (Carbide, Light Conversion). The laser wavelength was tuned to 1, 300 nm for three-photon excitation, generating a 433 nm THG signal. This THG signal was collected through a bandpass filter (FF02-447/60-25, Semrock). A homebuilt dispersion compensation unit ensured a pulse duration of approximately 60 fs at the sample plane. Depending on the skull thickness, 5 to 7.5 mw at the object was required to produce clear THG images.

### Microglia analysis

To quantitatively describe the morphological changes of microglia during the glaucoma, we measured the ramification index (RI) as previous work^41,61,62^. First, an imaging stack from Cx3cr1^GFP/+^ mice was taken. Next, a maximum intensity projection in the z direction was generated for every 20 µm. The threshold was then manually adjusted to cover the main branches and soma of the microglia, creating a binary mask. The microglia morphological changes were then quantified by custom-written Matlab code. The perimeter and area of the binary mask were measured, and the ramification index was calculated by:

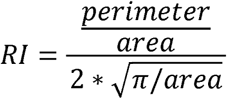

The surveillance index is calculated by simply measuring the area of the binary mask of each microglia, similar to previous work^63^.

To quantify the contrast of microglial processes, images obtained after two-photon ALPHA-FSS AO correction were used as a reference. A small region (∼10um* 10um) of microglial processes was manually selected from the maximum-intensity projection of AO-corrected images at different depths. Binary masks distinguishing the processes and background were generated using Otsu’s thresholding method in ImageJ. These masks were then applied to corresponding regions in both uncorrected and AO-corrected images to measure the average intensities of the processes (*I_p_*) and background (*I_b_*), enabling comparison of image contrast between blurred and corrected conditions. The contrast is defined as:

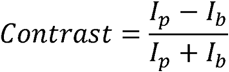

To quantify the relative fluorescence intensity of microglial processes and soma in images acquired without AO correction, binary masks of processes and soma generated from AO-corrected images were applied to the corresponding uncorrected images to measure the average fluorescence intensities. These values were then normalized to the corresponding intensities of processes and soma obtained from the AO-corrected images.

### Point spread function simulation

Electric field of aberrated 3D PSF is simulated based on the measured wavefront of wavefront sensor-based AO-TPEFM following the diffraction theory^64^. Two-photon PSF is calculated by the fourth power of the amplitude of electric field 3D PSF. The maximum intensity of two-photon PSF from an aberrated wavefront is normalized to that of a perfect wavefront. The integral value of aberrated two-photon PSF is calculated by adding the two- photon PSF intensity in a 15*15*15 *µm*^3^ volume, which is also then normalized with that of a perfect two-photon PSF.

### Imaging analysis

The images were processed using MATLAB or ImageJ. To eliminate inter-frame motion artifacts, the images were registered using the StackReg plugin in ImageJ, as well as the self-written algorithm in MATLAB.

### Statistical analysis

Statistical analysis and data visualization were performed using GraphPad Prism 10 software and MATLAB. The data are presented as mean ± SD, or ± SEM and α = 0.05 for all analyses. Data normality was assessed using the Shapiro–Wilk normality test. Normally distributed data were analyzed using paired, unpaired two-tailed t-test, one-way and two-way ANOVA test. Non-normally distributed data were analyzed using paired or unpaired non-parametric test (Wilcoxon test, Mixed-effects model analysis with Tukey’s multiple comparisons test). Some data were presented as contingency table. Fisher’s exact test is used for statistical analysis of contingency table. No statistical methods were used to determine the sample size.

## Declarations

## Ethics approval and consent to participate

All animal procedures in this study were conducted in accordance with the guidelines of the Animal Care Facility at the Hong Kong University of Science and Technology (HKUST) and were approved by the Animal Ethics Committee of HKUST.

## Consent for publication

All authors have consent to publish.

## Availability of data and materials

The datasets used and/or analyzed in the current research are available upon a reasonable request to the corresponding author.

## Authors’ contributions

Y.F., G.Y., Z.S. and J.Y.Q. conceived the research idea and designed the experiments; Y.F., Z.S. built and optimized the imaging systems; Y.F., G.Y. and K.L. performed animal surgery and imaging experiments; Y.F. and G.Y. analyzed the data with the help of Y.H.; Y.F. and G.Y. wrote the paper with inputs from all other authors.

## Acknowledgements

We thank Dr. Dan Zhu, Dr. Shaojun Liu from Huazhong University of Science and Technology, and Dr. Wei Feng, Dr. Chao Zhang from Zhanjiang Institute of Clinical Medicine for guidance in optical clearing window preparation, and Dr. Wenbiao Gan from Institute of Neurological and Psychiatric Disorders, Shenzhen Bay Laboratory, and Dr. Hang Zhou from Shenzhen University of Advanced Technology for sharing their expertise in thinned skull window preparation.

